# Detection of conspicuous and cryptic food by marmosets (*Callithrix jacchus*): An evaluation of the importance of color and shape cues

**DOI:** 10.1101/2021.04.11.439391

**Authors:** Priscilla Kelly Silva Barros, Felipe Nalon Castro, Daniel Marques Almeida Pessoa

## Abstract

Since 19th century, the adaptive function of color vision for object identification has fascinated science. Countless articles uncovered the advantages of trichromacy (i.e., color vision expressed by most humans) in detecting yellowish/reddish targets against a background of mature green leaves. Unfortunately, the same attention was not offered to achromatic visual information, that had their physiological foundations much more explored than their possible adaptive function. So far, mostly because of studies conducted in humans, we know that achromatic cues might also play an important role in object identification, particularly when camouflage is involved. For instance, dichromacy (i.e., color vision expressed by many colorblind humans), favors the detection of camouflaged targets by exploitation of shape cues. The present study sought to evaluate the relative importance of color and shape cues on the detection of food targets by female and male marmosets (*Callithrix jacchus*). Animals were observed with respect to their foraging behavior and the number of food targets captured. We confirmed that females are advantageous in detecting conspicuous food against a green background and revealed that females and males rely on shape cues to segregate cryptic food. Unexpectedly, males outperformed females in cryptic food foraging. Camouflage improved males’ (but not females’) performance.

**Highlights:**

- Shape cues alone improve capture of cryptic targets by female and male marmosets.
- The sole use of color cues leads to clear-cut foraging performances between sex.
- Females outperform males when searching for orange food against a green background.
- Males outperform females when searching for green food against a green background.
- Males match females’ performance when searching for orange camouflaged food.

## 1. INTRODUCTION

For an animal to have color vision, it must possess, at least, two distinct classes of photoreceptor cells in its retina, in addition to a neural network capable of comparing the responses of these cells (Cowey and Heywood 1995). Literature has shown that the greater the variety of retinal photoreceptor cells, the better the animal’s ability to distinguish color (Bowmaker 1998). In this context, diurnal mammals exhibit dichromatic color vision (Jacobs 2018), associated with two photopigment classes, as observed in some colorblind humans. Yet, primates (Jacobs 2010) appear to have evolved three different types of cones in their retina. This results in trichromatic vision, like that found in humans with normal color vision.

The distribution of color vision among primates has a peculiar pattern. Uniform trichromacy is observed among Old World monkeys and apes (Catarrhini), in which most individuals within a population exhibit three different classes of cone photopigments: S pigments, which preferentially absorb short wavelengths (blue range); M pigments, which preferentially absorb medium wavelengths (green range); and L pigments, which preferentially absorb long wavelengths (red range). While the gene encoding the S pigment is found on an autosomal chromosome, M and L pigments are coded on the X chromosome (Jacobs 2010).

Alternatively, New World primates (Platyrrhini) display a visual polymorphism, characterized by the occurrence of females with trichromatic (heterozygosis) or dichromatic (homozygosis) vision, and obligatory dichromatic males (hemizygosis) (Hunt et al. 1993). This visual polymorphism occurs because a single gene encodes M/L pigments, and is located on the X chromosome, in addition to a gene that encodes the S pigment on an autosomal chromosome (Carvalho et al. 2017). Different visual alleles can be found on the X chromosome, which produce photopigments with distinct light absorption peaks, yielding trichromats with S, M and L cone pigments, and dichromats with S and only one M or L pigment.

On one hand, color is considered an important visual cue, since it facilitates the identification of objects and patterns within a scenario, as well as assisting in memorization (Wichmann et al. 2002). The visual recognition of objects can be improved by simply adding color to an image (Shevell and Kingdom 2008). Since the 19^th^ century, when the naturalist Grant Allen proposed, in 1879, that the evolution of fruit coloration and animal color vision had occurred in parallel, color vision in primates has been understood as resulting from an adaptive process involving the identification of conspicuously colored food resources (see Jacobs 2007). In line with that, the frugivory and folivory hypotheses were postulated to explain which advantages trichromats would have in detecting ripe fruit and young leaves, respectively, against a background of mature green leaves (Mollon 1989; Sumner and Mollon 2000; Dominy and Lucas 2001).

On the other hand, by using color to camouflage objects, that is, adding chromatic noise, one can hamper object identification, especially when other cues (e.g., size, shape, texture) are not available (Liebe et al. 2009). This happens because neural processing of chromatic and achromatic information occurs concurrently and competitively. Thus, as argued by Regan and collaborators (2001), primates with greater dependence on color cues (e.g., trichromats) should be more affected by chromatic noise, that is, they would focus a significant amount of their neural processing to deal with chromatic information, at the cost of a worse processing of achromatic cues. Alternatively, those primates with poorer color vision (e.g., dichromats) would benefit from a reduced chromatic noise, being able to redirect additional neural processing to achromatic information assessment, important for the identification of shapes, outlines, and textures. In fact, Mollon (1989) predicted that, given their possible disadvantage in detecting conspicuous food items, dichromats should compensate their inferior performance by identifying camouflaged objects. Investigations presenting camouflaged geometric figures to humans (Morgan et al. 1992; Saito et al. 2006) and non-human primates (Saito et al. 2005) showed that dichromats, when compared with trichromats, were more capable of breaking camouflage and detected hidden shapes. Indeed, some naturalistic/ecological studies gave support to the idea that the visual polymorphism of Neotropical primates balances (Hiwatashi et al. 2010) the advantages of dichromats in recognizing hidden objects in the environment (Caine et al. 2003; Melin et al. 2007; Smith et al. 2012) and those of trichromats in detecting conspicuous targets (Caine and Mundy 2000; Smith et al. 2003; Melin et al. 2013, 2017; Pessoa et al. 2014; Abreu et al. 2019).

Research on *Branisella boliviana*, a 26 million-year-old Neotropical primate fossil, weighing approximately 760 grams (Kay et al. 2002) and, possibly, exhibiting visual polymorphism (Heesy and Ross 2001), suggests that leaves had never been a significant component of the diet of Neotropical primate ancestors, which likely fed on fruits and insects (Kay 1984). This leads to the inference that folivory has never been important in the evolution of callitrichids (i.e., marmosets and tamarins), which are the smallest extant Neotropical primates (weighting less than 800g; Arruda et al. 2019). In addition, it has been proposed that insectivory and predation pressure are selective factors that maintained the visual polymorphism in this taxon (Pessoa et al. 2014; De Moraes et al. 2021).

In the past decades, color vision of common marmosets (*Callithirx jacchus*) has been substantially examined through theoretical, behavioral, electrophysiological and molecular approaches (Travis et al. 1988; Tovée et al. 1992; Hunt et al. 1993; Wilder et al. 1996; Shyue et al. 1998; Kawamura et al. 2001; Surridge and Mundy 2002; Pessoa et al. 2005a; Freitag and Pessoa 2012; Pessoa et al. 2014; Moreira et al. 2015; Abreu et al. 2019; Pessoa and Freitag 2019; Mantovani et al. 2020), showing that *C. jacchus* expresses a tri-allelic visual polymorphism, that encodes M/L pigments with maximum absorption at 543 nm, 556 nm and 562 nm, and allows the existence of six visual phenotypes (three dichromats and three trichromats) in natural populations (Jacobs 2007).

Although many naturalistic studies have already addressed the possible advantages and disadvantages of color vision in food identification (Caine and Mundy 2000; Caine et al. 2003; Smith et al. 2003; Leonhardt et al. 2009; Freitag and Pessoa, 2012; Smith et al. 2012; Bompas et al. 2013; Melin et al. 2013; Pessoa and Freitag, 2019), we believe the relative importance of different visual information (e.g., shape and color) in food target detection has not been sufficiently analyzed in non-human primates. This is due because the attention of previous studies has fallen almost exclusively on controlling stimuli’s color, while shape has been overlooked. As such, the present study sought to examine the influence of color and shape cues on food detection by female and male marmosets (*Callithrix jacchus*). Here, for practical reasons, and since, at least, two thirds of females are expected to be trichromats (Jacobs 2007), while the totality of males are dichromats, we assume that females, as a group, will behave as trichromats, while males will perform as dichromats, an approach that has been successfully adopted by behavioral ecologists when visual phenotypes are unknown (Dominy et al. 2003, Yamashita et al. 2005). We hypothesize that chromatic (i.e., color) and achromatic (i.e., shape) signals, as well as chromatic and achromatic noise, will influence food detection by female and male marmosets. We also theorize that animals of different sex will display contrasting performances under conditions in which color cues play a decisive role in food advertisement. Our predictions are that, when foraging on a background of dappled green cubical elements (i.e., in the presence of lower chromatic noise and higher achromatic noise): (1) females will outperform males when searching for orange cubical food items (i.e., when color cues are available only for trichromatic animals), but not for green (cryptic condition) or blue (i.e., when color cues are available for trichromats and dichromats) cubical targets; (2) females and males will segregate green spherical targets (i.e., when shape cues are available for all subjects) much easier than green cubical ones (cryptic condition); and (3) females will outperform males when searching for orange spherical items (i.e., when color and shape cues are available for trichromats, while only shape cues are available for dichromats). Otherwise, when foraging on a background of dappled green and orange cubical elements (i.e., in the presence of higher chromatic noise and lower achromatic noise): (4) males will outperform females when searching for orange spherical items (i.e., when shape cues will be available for dichromats, while will be masked for trichromats by chromatic noise).

## 2. MATERIAL AND METHODS

### 2.1. Subjects

From a breeding colony of, approximately, 200 common marmosets (*Callithrix jacchus*), maintained by the Federal University of Rio Grande do Norte’s Primate Center, we selected five family groups to take part in the experiments. Females and males of different age classes were monitored, but only data from adults, subadults and juveniles were analyzed in the present study, totaling 10 females and 15 males. All behavioral experiments were conducted in 2010 so, since we did not collect tissue samples from our subjects by then, determination of individual color vision phenotypes was not possible.

To avoid the stress of daily capture and transportation to a new environment, family group had access to experimental enclosures (1.0 m width x 2.0 m length x 2.0 m height), containing platforms and branches of different heights, that was adjacent to their larger home enclosures (2.0 m width x 2.0 m length x 2.0 m height). Temperature and sunlight varied freely, according to local weather. Experimental sessions were always conducted in the mornings (07:00 a.m. to 10:00 a.m.), when food trays were removed from the enclosures. Water was always provided *ad libitum*. The experimental procedures were approved by the Animal Research Ethics Committee of the Federal University of Rio Grande do Norte (Proc.045/2009) and adhered to the legal requirements of Brazilian regulation. Experiments complied with the ARRIVE guidelines and were carried out in accordance with the National Institutes of Health guide for the care and use of Laboratory animals (NIH Publications No. 8023, revised 1978).

### 2.2. Stimuli

We produced visual targets using cake frosting (Arcolor), with the following composition: sugar, vegetable fat, vegetable gums, and glucose syrup. Food targets were molded into spheres or cubes (approximately 1 centimeter wide) and colored with food dyes (Arcolor), consisting of glucose syrup, modified starch, humectant propylene glycol, water, preservatives, and organic dyes. For each target hue (i.e., blue, green, or orange) we produced two variants with different brightness levels (e.g., light, and dark blue targets). We also produced background elements from white foam rubber sheets (EVA - Ethylene Vinyl Acetate) that were cut into small cubes (about 1 cm^3^), divided into five similar piles, and coated with different water-based paints, creating four different shades of green and one shade of orange. Once finished, the surface of food targets and background elements appeared matte and polished. Using food targets and background elements of the same size, same texture and variable brightness levels was a strategy adopted by us to preclude the identification of food targets based on visual cues other than color and/or shape.

Fine adjustments in stimuli colors were performed by a human trichromat (P.K.S.B.) and a human dichromat (D.M.A.P.), to ensure that, on a green dappled background, green-colored targets were cryptic and blue-colored targets were conspicuous. These human subjects also ensured that orange-colored targets had the same appearance as orange-colored background elements. Orange-colored targets were also adjusted to seem conspicuous to trichromats (P.K.S.B.) and cryptic to dichromats (D.M.A.P.), when viewed against a green dappled background.

### 2.3. Spectral Measurements

We measured stimuli’s coloration with a portable spectrometer (USB4000 UV-VIS Fiber Optic Spectrometer, Ocean Optics Inc.) running Spectra Suite software (Ocean Optics Inc.), connected to a bifurcated optical fiber (R400-7-VIS-NIR, Ocean Optics Inc.), that was attached to a probe-holder (RPH-1, Ocean Optics Inc.). The system was illuminated with an artificial light source (LS-1, Ocean Optics, Inc.). For system calibration we measured light reflecting from a white standard surface (WS-1, Ocean Optics Inc.), and obstructed light coming from the light source for determining the black standard. After calibration, we measured the reflectance spectra (coloration) of food targets and background elements, positioning the tip of the optical fiber 0.5 cm from the measured surface, always at an angle of 45°. The number of spectra averaged to scan, and boxcar were always set to 10 and 5, respectively. For each kind of stimuli, we randomly picked four samples, measured their coloration, and generated an average reflectance spectra curve (Figure 1).

**Figure 1.**
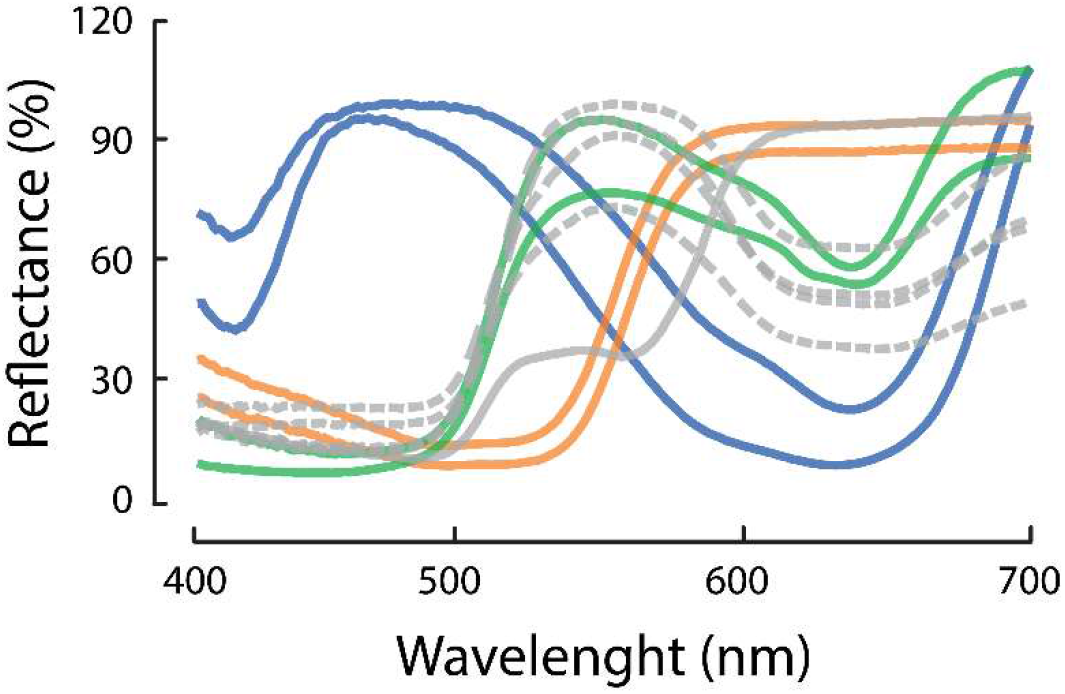
Stimuli reflectance spectra. Gray dashed lines represent the coloration of green background elements (four different shades are shown), while the gray solid line illustrates the reflectance of orange background elements. Solid colored lines represent food targets coloration, in two different shades of blue (curves peaking around 450 nm), orange (curves peaking around 600 nm) and green (curves peaking around 550 nm).

### 2.4. Visual Modeling

Visual modeling was conducted in *pavo 2*.*0* (Maia et al. 2019), a R package for spectral analysis of color (R Core Team 2020). We started estimating the absolute amount of light captured by marmosets’ photoreceptors (‘Qi’ – quantum catch), considering three factors: a) the reflectance spectrum of each stimulus (Figure 1); b) the illuminant spectrum of enclosures’ ambient light; c) and marmoset’s visual sensitivity curves (for further details see Perini et al. 2009). Then, we set the remaining parameters to default (i.e., trans = ‘ideal’, vonkries = FALSE, scale = 1), and set relative = FALSE, since we required absolute quantum catches, instead of relative quantum catches.

By generating MacLeod–Boynton chromaticity/luminance diagrams, which graphically represent how stimuli should excite marmosets’ red-green, blue-yellow and luminance visual channels, we modeled how trichromatic and dichromatic marmosets would discriminate food targets from background elements. We constructed the diagrams based on absolute quantum catch information for S (Q_S_), M (Q_M_) and L (Q_L_) photopigments. Since marmosets are polymorphic, and express three kinds of trichromatic phenotypes (with S/M/L spectral peaks at 430/543/556 nm, 430/543/562 nm, and 430/556/562 nm) and three kinds of dichromatic phenotypes (with S/L spectral peaks at 430/543 nm, 430/556 nm, and 430/562 nm), we modeled chromaticity/luminance diagrams for six different visual phenotypes. While trichromats exhibit three visual channels, Q_L_/(Q_M_+Q_L_) (i.e., red-green channel), Q_S_/(Q_M_+Q_L_) (i.e., blue-yellow channel), and Q_M_+Q_L_ (i.e., luminance channel), dichromats only display two, a blue-yellow (Q_S_/Q_L_) and a luminance channel (Q_L_).

### 2.5. Experimental procedure and behavioral observations

We adjusted the number of food targets offered according to family size (disregarding infants), preserving a ratio of two targets per animal (e.g., 10 targets were offered to a family composed of 5 animals). Then, we randomly scattered cubical or spherical targets among three, out of six, plastic trays (each measuring 30 cm width x 50 cm length x 9 cm height), previously filled with green (lower chromatic noise conditions) or green and orange (higher chromatic noise condition) background elements, which we placed on the floor of the experimental enclosures (Figure 2).

**Figure 2.**
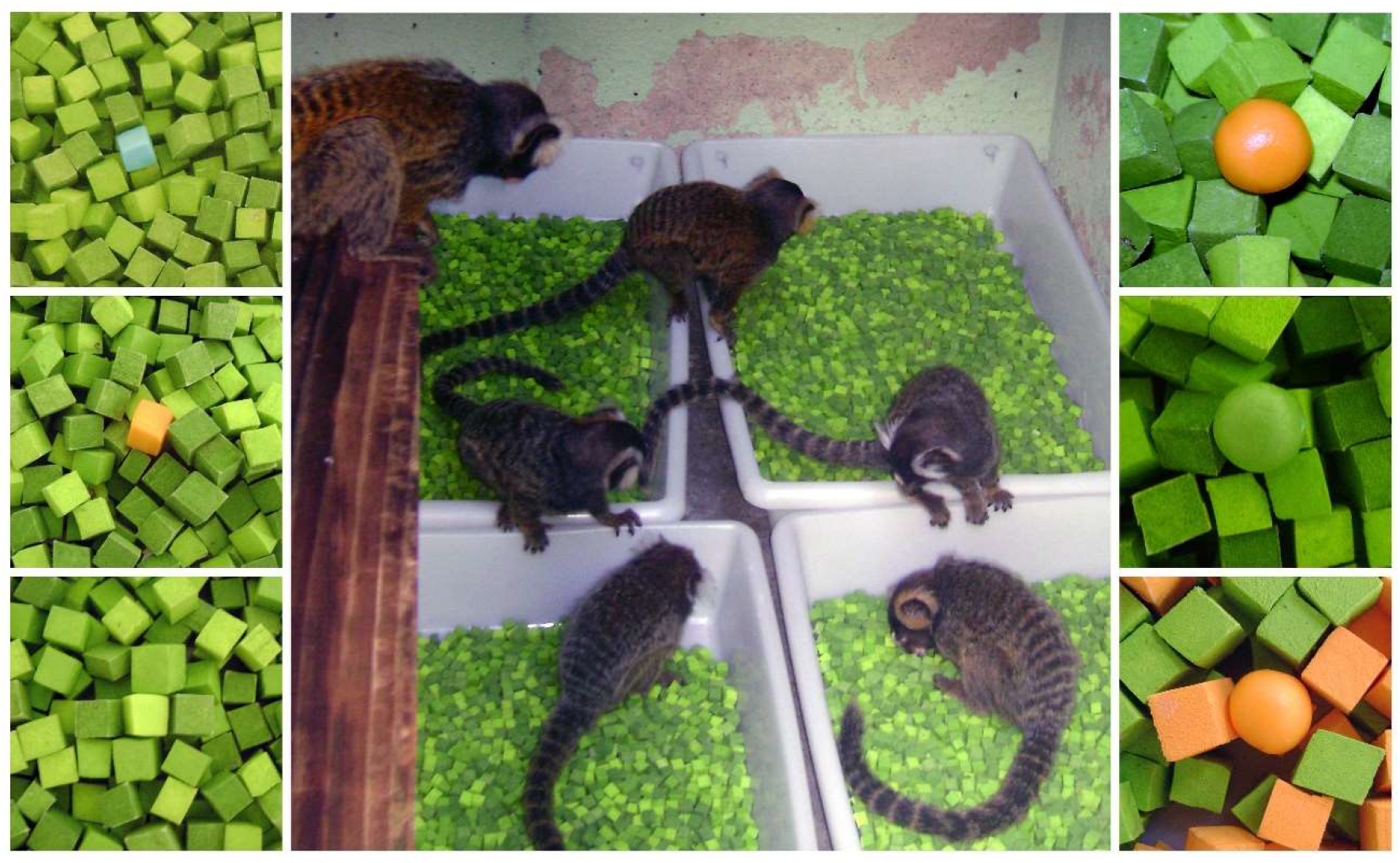
Subjects engaged in foraging activity (center photograph). Lateral panels illustrate six different setups used in the experiments. Upper left: green background elements with a blue cubical-shaped food target. Middle left: green background elements with an orange cubical-shaped food target. Bottom left: green background elements with a green cubical-shaped food target. Upper right: green background elements with an orange spherical-shaped food target. Middle right: green background elements with a green spherical-shaped food target. Bottom right: green/orange background elements with an orange spherical-shaped food target.

To each marmoset family group, we repeatedly presented (22 repetitions) six experimental setups (Figure 2): (A) Four shades of green background elements with two shades of blue cubical-shaped food targets; (B) Four shades of green background elements with two shades of orange cubical-shaped food targets; (C) Four shades of green background elements with two shades of green cubical-shaped food targets; (D) Four shades of green background elements with two shades of orange spherical-shaped food targets; (E) Four shades of green background elements with two shades of green spherical-shaped food targets; (F) Two shades of green and one shade of orange background elements with two shades of orange spherical-shaped food targets.

Each experimental session, lasting 10 minutes, began with one of the human observers opening a wooden sliding door and allowing the subjects into the experimentation enclosure. Presentation order for the six setups was randomized, as was the time of the day for observing each family. On each experimental day, we exposed each family to the same experimental setup, while pseudo-randomly selecting a focal individual from each family (Altmann 1974) to be monitored regarding its foraging behavior. Foraging behavior was defined as the visual scanning behavior directed towards the plastic trays, as well as the behavior of actively searching (i.e., while on the plastic trays, slowly walking and looking towards the stimuli) and capturing food targets. We did not compute food manipulation and/or intake as foraging. Alongside, a second human observer was responsible for using all-occurrence sampling (Altmann 1974) to record the number of targets captured by every individual in the group.

We left infants out of the analysis since their behaviors were not registered. We were unable to register the foraging behavior (focal sessions) of one of the juvenile males, although we recorded the number of targets captured by it in every experimental session. In addition, after we had already finished monitoring the foraging behavior (focal sessions) of one of the adult females, it fell ill and deceased, so the number of targets captured by it could not be considered in our analysis. Except for these two incidents, every animal that managed to capture a food item was registered with respect to food acquisition. So, at the end of the study period, four complete focal sessions (measuring foraging duration) had been monitored for each adult and subadult marmoset (ten females and fourteen males), in a total of 16 hours of sampling effort (40 minutes for each focal subject). In addition, for each family group, 22 sessions determining the number of targets captured were accomplished and included data from nine females and 15 males, in a total of 220 minutes of sampling effort. We used two chronometers and two audio recorders to register foraging information.

### 2.6. Statistical Analysis

The performance of test subjects was analyzed according to their average foraging time and average percentage of food targets captured. For calculating average foraging time, we considered the total amount of time (in seconds) spent in foraging (while food targets were still available) by each animal, dividing it by total session duration, which ended when all food targets were consumed or after 10 minutes had passed. The average percentage of food targets captured was determined considering the ratio between total number of targets captured per animal and total number of targets offered for that specific family group.

We ran six independent General Linear Models (GLM), with post-hoc Bonferroni, for testing the effect of target shape, target color and background chromatic noise for each of the dependent variables (i.e., average foraging time and average percentage of food targets captured). We evaluated the effect of target shape by comparing experimental setups (C) and (E), assigning target shape (round or square) and sex (female or male) as independent variables. To explore the effect of target color, we compared experimental setups (A), (B) and (C), selecting target color (orange, green or blue) and sex (female or male) as independent variables. Finally, we explored the effect of background chromatic noise by comparing experimental setups (D) and (F), defining background color (green or orange-green) and sex (female or male) as independent variables. In all analyzes, the alpha considered was 5%.

## 3. RESULTS

### 3.1. Visual modeling - Chromaticity/luminance diagrams

In chromaticity/luminance diagrams, stimuli similarity is given as a measure of spatial proximity, i.e., the further apart two stimuli are represented, the more dissimilar they are, and vice-versa. With that said, Figures 3 and 4 predict, on one hand, that green targets should be cryptic to dichromatic and trichromatic marmosets, since they overlap with the green background elements with respect to chromaticity and luminance. But, on the other hand, blue targets must be highly conspicuous against green background elements to dichromatic and trichromatic individuals, when considering chromaticity, but not luminance. Figures 3 and 4 also predict that orange food targets should be cryptic to dichromats and conspicuous to trichromats, since their chromaticity overlap with green background elements on the blue-yellow channel, but not on the red-green visual channel. In addition, compared to orange background elements, orange targets should always be cryptic.

**Figure 3.**
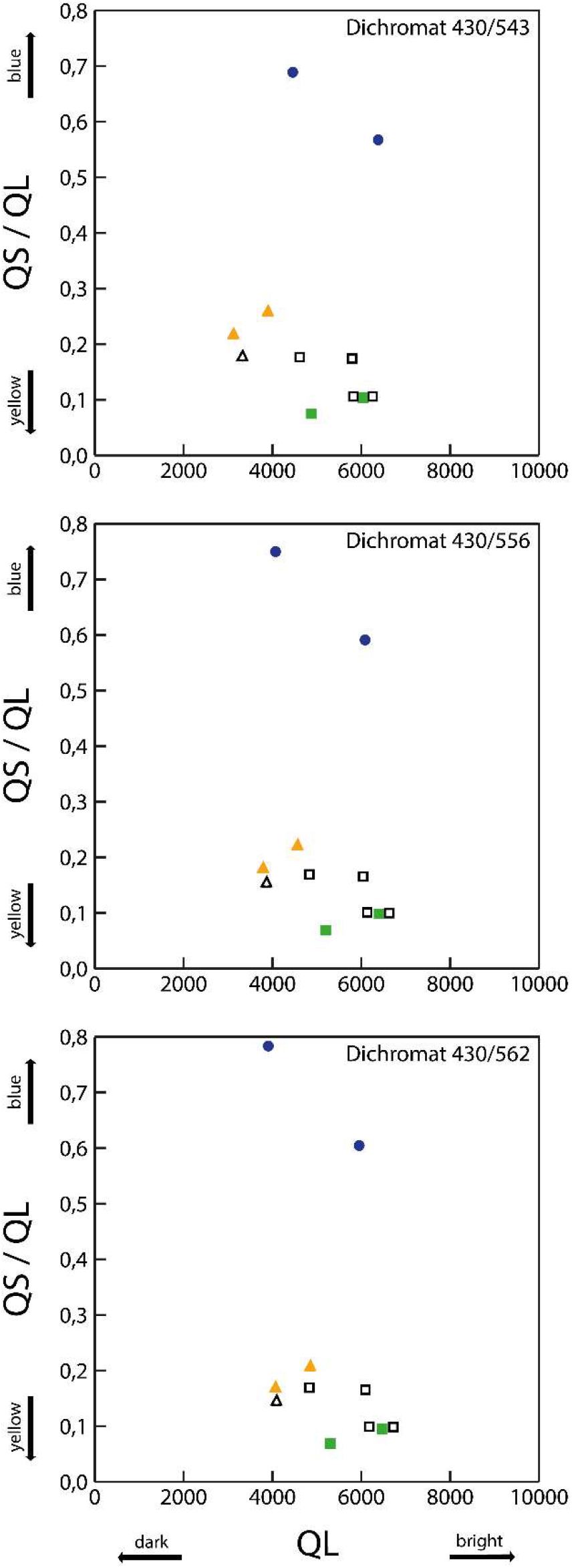
Chromaticity/luminance diagrams for three dichromatic phenotypes found in marmosets. Blue (closed circles), orange (closed triangles) and green food targets (closed squares), as well as orange (open triangles) and green (open squares) background elements, are plotted according to how they activate the chromatic blue-yellow (QS/QL) axis and the luminance (QL) axis of marmoset dichromatic vision.

**Figure 4.**
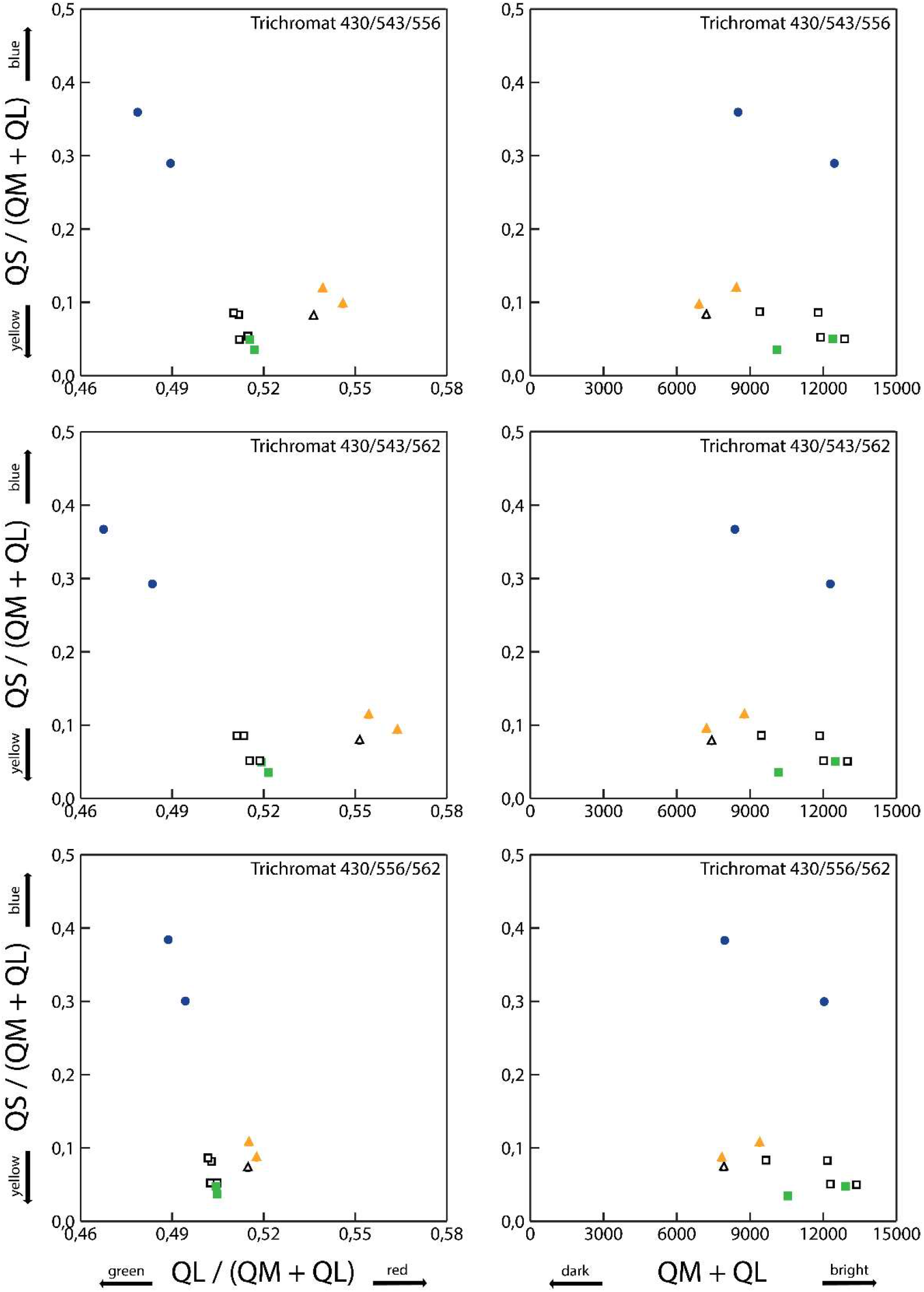
Chromaticity/luminance diagrams for three trichromatic phenotypes found in marmosets. Blue (closed circles), orange (closed triangles) and green food targets (closed squares), as well as orange (open triangles) and green (open squares) background elements, are plotted according to how they activate the chromatic red-green [QL/(QM+QL)] and blue-yellow [QS/(QM+QL)] axes, and the luminance (QM+QL) axis of marmoset trichromatic vision.

### 3.2. Influence of target color

Target color influenced foraging time (F_2,42_=16.90; p<.001; η^2^_Partial_=0.446) and percentage of targets captured (F_2,44_=3.88; p=.028; η^2^_Partial_=0.150). On one hand, both female and male marmosets significantly invested more time foraging for green food, when compared to their time spent looking for orange or blue items (Figure 5B). On the other hand, females captured fewer green food items than orange ones, while males had a bad time capturing orange food, which were significantly less captured than green and blue targets, and a good time capturing blue targets, which were taken more frequently than green and orange targets (Figure 5A).

**Figure 5.**
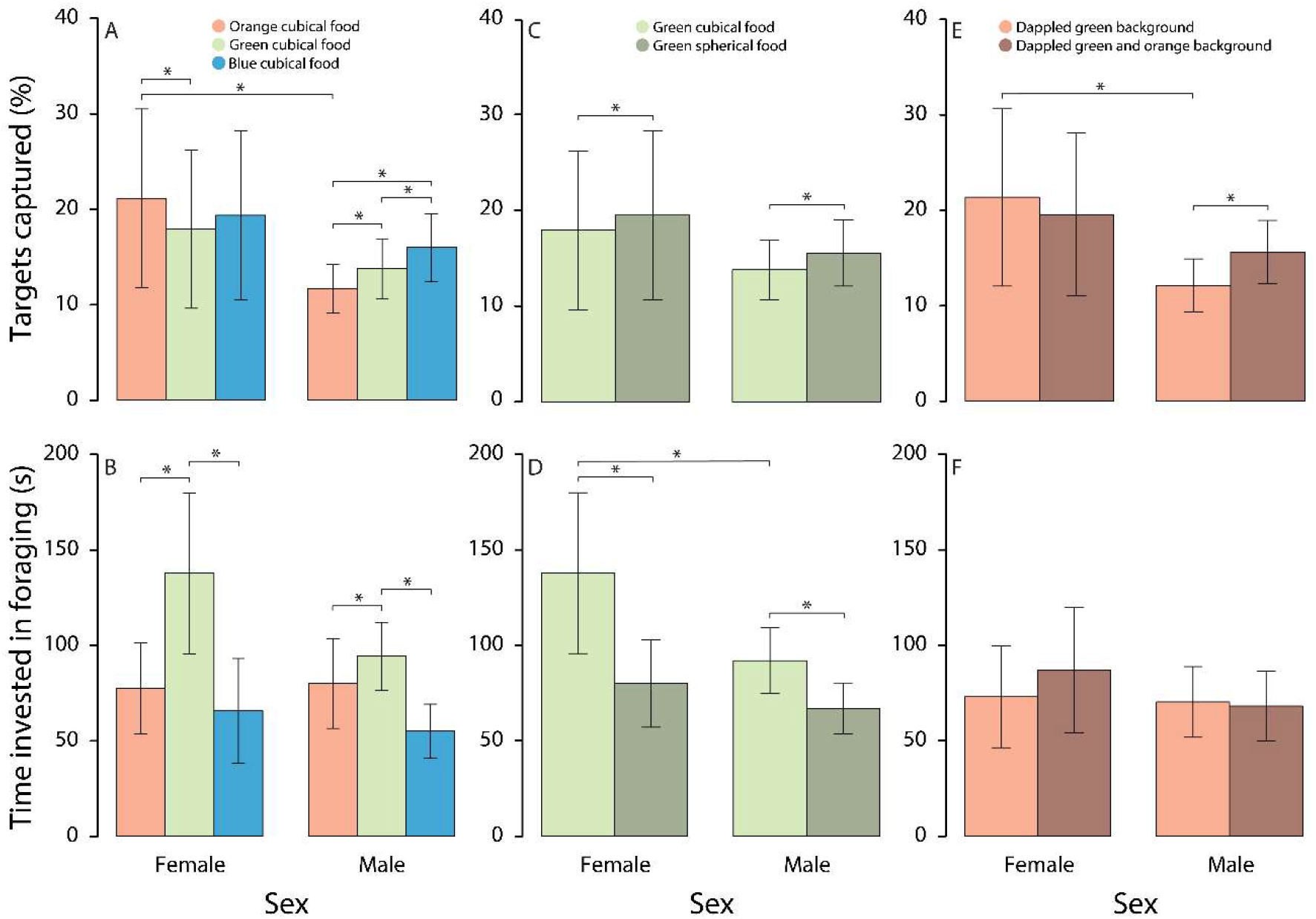
Influence of target color, target shape and background chromatic noise on the feeding performance (average foraging time and average percentage of targets captured) of female and male captive marmosets (*Callithrix jacchus*). Confidence intervals are represented by vertical whiskers. Horizontal bars with asterisks indicate significant differences for α=.05.

We also found an interaction between target color and sex (F_2,44_=12.25; p<.001; η^2^_Partial_=0.358) for percentage of targets captured, with females capturing significantly more orange food than males (Figure 5A).

### 3.3. Influence of target shape

Shape of food items significantly influenced foraging time (F_1,22_=34.20; p<.001; η^2^_Partial_=0.609) and percentage of targets captured (F_1,22_=9.60; p=.005; η^2^_Partial_=0.304), favoring the detection of spherical over cubical targets (Figures 5C and 5D).

We also found an interaction between target shape and sex (F_1,22_=5.40; p=.030; η^2^_Partial_=0.197) regarding foraging time, with females investing more time foraging for green cubical food than males (Figure 5D).

### 3.4. Influence of background chromatic noise

Background chromatic noise had no influence on foraging time invested by females or males (F_1,22_=0.55; p=.463; η^2^_Partial_=0.025). Yet, concerning the percentage of targets captured, we found an interaction between background chromatic noise and sex (F_1,22_=11.50; p=.003; η^2^_Partial_=0.343). On one hand, females captured orange spherical targets on a dappled green background more frequently than males. On another hand, males significantly captured more orange spherical food items deposited on a dappled green and orange background than on a dappled green background (Figure 5E).

## 4. DISCUSSION

### 4.1. Influence of target color

Our behavioral results show a clear-cut difference between female and male subjects’ performance and are in line with data modeled by our chromaticity/luminance diagrams, corroborating our first prediction that females should outperform males when searching for orange cubical targets, but not for green or blue cubical food. These results are a clear indication that our female subjects, as a group, indeed behaved as trichromatic individuals, outperforming dichromatic marmoset males in red-green discrimination tasks, as extensively documented by literature for captive marmosets (Caine and Mundy 2000; Caine et al. 2003; Pessoa et al. 2005b). Although our study subjects were not genotyped with respect to their color vision, we believe differences between female and male performance shown here cannot be chiefly attributed to innate sex differences, instead of color vision phenotype differences, since no difference in female and male performance were found when animals were searching for blue food items (i.e., the positive control condition). In addition, It is important to highlight that, although our chromaticity/luminance diagrams sometimes depict food targets as being darker or lighter than background elements (i.e., overlapping is not perfect), the use of brightness cues by our experimental subjects was almost certainly prevented, once we varied the brightness of our targets and background elements, making luminance cues unreliable.

### 4.2. Influence of target shape

Regarding the influence of shape cues on target detection, we confirmed our second prediction that females and males would benefit from shape cues and segregate green spherical targets much easier than green cubical ones. Unexpectedly, we also found that dichromatic males invested a lower amount of time searching for cryptic targets (i.e., green cubical food), outperforming females. This dichromatic advantage for cryptic food detection contradicts what has been found by psychophysical experiments, that presented human subjects with photos of wild fruits (Melin et al. 2013) and revealed a trichromatic advantage for cryptic fruit detection. Also contradicts a behavioral ecology study conducted in tamarins (Smith et al. 2012), which concluded that trichromats were superior to dichromats at detecting green generalist insects. Furthermore, previous naturalistic studies involving captive marmosets (Caine and Mundy 2000; Caine et al. 2003) have failed to show any significant dichromatic advantage for cryptic food identification.

Despite their known importance for object recognition (Liebe et al. 2009), shape cues had not yet been controlled and evaluated in naturalistic experiments examining foraging of colored food items by non-human primates. In theory, regardless of information concerning color, shape cues may be capable of indicating the presence of food (e.g., fruits and insects) in vegetation. When objects are conspicuously colored, the relative importance of shape may decline. However, in the presence of chromatic noise (e.g., disruptive coloration camouflage) or in situations in which coloration conceals targets (e.g., background matching camouflage), shape information should become more significant (Saito et al. 2005). Our results confirm that shape cues are important under seminatural conditions and play a complementary role in food identification. In fact, according to Párraga and collaborators (2002), the shape of fruits closely match those that our visual system finds optimal. In the present study, when we tested the effect of shape cues on food identification, the influence of other visual cues (e.g., color, brightness, texture, gloss, and size) was considered negligible, once they were either controlled (i.e., stimuli’s color, size, texture, and gloss) or randomized (i.e., stimuli’s brightness).

### 4.3. Influence of background chromatic and achromatic noise

Presenting a performance consistent with that of trichromats, female marmosets outperformed dichromatic males in tasks involving the detection of orange spherical targets against a dappled green background, confirming our third prediction. However, our data also rejected our fourth prediction that males would outperform females when searching for orange spherical items under camouflaged conditions. Instead, we revealed that camouflage did not influence the performance of female subjects, that captured comparable amounts of orange spherical food against both kinds of backgrounds tested in the present study. Still, in contrast, camouflage exerted a significant impact on male feeding success. Under the camouflaged condition (i.e., dappled green and orange background) dichromatic males improved their detection of orange spherical targets, when compared to the non-camouflaged condition (i.e., dappled green background), matching females’ performance. The influence of a camouflaged background on conspicuous food detection by dichromatic marmosets contradicts what have been presented by a previous study (Caine et al. 2003). Caine and collaborators (2003) found that camouflaged, in comparison to non-camouflaged conditions, significantly reduces feeding performance in trichromatic marmosets, but not dichromatic ones. We believe this discrepancy could be attributable to differences in the spectral features of the background elements used in each study. According to the reflectance spectra of background elements, it appears that Caine’s non-camouflaged green background was less variegated (i.e., presented a reduced variance in brightness) when compared to ours, what might have generated lower levels of achromatic noise. Alternatively, our dappled green background must have produced much more achromatic noise, which possibly made tasks more difficult and resemblant to natural challenges.

Both of our unexpected results, the dichromatic advantage for cryptic food identification and the dichromatic improvement in conspicuous food detection under camouflaged conditions might be explained by how primate visual systems are organized. In primates, hierarchical processing of chromatic and achromatic information flows in parallel pathways and tend to compete for computational power, in such a way that color processing hampers achromatic processing (including shape information) and vice-versa (Tovée 2008). Consequently, an animal devoid of color vision, or with a reduced color discrimination capability (e.g., dichromat), would reserve most of the processing power allocated to visual processing to its parallel achromatic pathway, improving the detection of contours, textures, and shapes (Morgan et al. 1992; Saito et al. 2005, 2006). Alternatively, in trichromats, an additional color processing pathway (red-green color opponency), absent in dichromats (Mollon 1989), would demand processing power that would have to be redirected from the achromatic pathway.

Background coloration exerts a substantial influence on object identification, given that it can increase target segregation based on their color contrast, as well as promote perceptual confusion, by modifying the color of objects (Shevell and Kingdom 2008) and introducing chromatic and/or achromatic noise (Liebe et al. 2009). Since it supposedly presents less achromatic noise then the dappled green background, the dappled green and orange background must have facilitated the detection of spherical orange targets by dichromatic males based on shape cues, which explains the improvement in male’s performance under camouflage conditions. The dichromatic visual system of male marmosets cannot segregate orange and green background elements, so males might have perceived the camouflaged background as roughly homogenous, barely being affected by its chromatic noise, what would allow these dichromatic individuals to effortlessly use the contour of orange spherical targets to segregate them from the cubical background elements. In contrast, the same condition posed a problem to females because, for trichromats, the dappled green and orange background must have significantly enhanced chromatic noise, disturbing shape perception.

At first, it might seem unexpected that male dichromats outperformed females when foraging for cryptic food (i.e., green cubical food) under a highly achromatically noisy condition (i.e., dappled green background), since achromatic noise should have adversely affected both male and female performances. Once, due to its four different shades (brightness) of green background elements, the non-camouflaged background must have appeared extremely heterogeneous to all animals (Párraga et al. 2002). One possible explanation to dichromatic advantage would be that, even though green targets look chromatically cryptic when compared to green background elements, discrete red-green chromatic differences between targets and background elements could have generated enough chromatic noise to affect the discrimination of food by trichromats. Chromatic noise generated by the blue-yellow chromaticity axes of dichromats and trichromats, in turn, would be a shared disadvantage, so it would not be significant for their differential performance.

## 5. CONCLUSION

Here, we confirmed that trichromats are advantageous in orange food detection. Surprisingly, we unveiled a priviously unknown dichromatic advantage for green food identification and found that background color impacts target identification by dichromatic marmosets, but not trichromats. These findings reinforce the idea that visual polymorphism in Neotropical primates is maintained through balancing selection (Hiwatashi et al. 2010), and it would involve multiple advantages and disadvantages of dichromats and trichromats (Pessoa et al. 2014).

Although blue fruits have been reported to comprise the diet of some primate species (Savage et al. 1987; Dominy 2004; Perini et al. 2009), this is only the second Neotropical primate naturalistic study (besides Freitag and Pessoa 2012) to explore the detection of blue food items against a green dappled background. Our data have demonstrated that female and male captive marmosets have no trouble identifying blue food items against a dappled green background.

In summary, our results show that both color and shape cues are important sources of visual information that influence target discrimination according to the degree of prevailing background noise. However, it is essential to remember that color vision is a complex phenomenon that also depends on other object features and on environmental luminosity, such as stimuli size (Gomes et al. 2005) and ambient light intensity (Pessoa and Freitag 2019). We urge future naturalistic studies to investigate underexplored visual factors such as stimuli’s brightness (Hiramatsu et al. 2008), texture, gloss, and size (Valenta et al. 2020).

## 6. CRediT AUTHOR STATEMENT

**Priscilla K.S. Barros**: Conceptualization, Methodology, Validation, Investigation, Data curation, Writing - Original Draft, Writing - Review & Editing, Project Administration. **Felipe N. Castro**: Formal analysis, Data curation, Writing - Review & Editing. **Daniel M.A. Pessoa**: Conceptualization, Methodology, Resources, Writing - Review & Editing, Visualization, Supervision, Funding acquisition.

## 7. ACKNOWLEDGEMENTS

This study was financed in part by the Coordenacao de Aperfeicoamento de Pessoal de Nivel Superior – Brazil (CAPES), Finance Codes 001 and 043/2012, that also provided a M.Sc. Scholarship to P.K.S.B.; by Conselho Nacional de Desenvolvimento Cientifico e Tecnologico – Brazil (CNPQ), Finance Codes 478222/2006-8 and 474392/2013-9, that also provided a Researcher Scholarship to D.M.A.P.; and by Programa de Apoio aos Nucleos de Excelencia – FAPERN/CNPq, Finance Code 25674/2009. The funding sources were not involved in study design, collection, analysis, interpretation of data, writing or in the decision to submit the article for publication. We would like to thank the Primate Center’s staff for animal care and maintenance, specially J.F.V. Coutinho and E.C. Rodrigues. We are also indebted to L.E.F. Carvalho, S.R.N. Silva and S.S. Farias for assisting in data collection.

